# Unraveling Alzheimer’s Disease: Investigating Dynamic Functional Connectivity in the Default Mode Network through DCC-GARCH Modeling

**DOI:** 10.1101/2024.06.02.597071

**Authors:** Kun Yue, Jason Webster, Thomas Grabowski, Hesamoddin Jahanian, Ali Shojaie

**Affiliations:** Department of Biostatistics, University of Washington, Seattle; Department of Radiology, University of Washington, Seattle; Department of Neurology, University of Washington, Seattle

**Keywords:** Dynamic Brain Functional Connectivity, Alzheimer’s Disease, Default Mode Network, Resting-state fMRI, Amyloid Beta Biomarker

## Abstract

Alzheimer’s disease (AD) has a prolonged latent phase. Sensitive biomarkers of amyloid beta (*Aβ*), in the absence of clinical symptoms, offer opportunities for early detection and identification of patients at risk. Current *Aβ* biomarkers, such as CSF and PET biomarkers, are effective but face practical limitations due to high cost and limited availability. Recent blood plasma biomarkers, though accessible, still incur high costs and lack physiological significance in the Alzheimer’s process. This study explores the potential of brain functional connectivity (FC) alterations associated with AD pathology as a non-invasive avenue for *Aβ* detection. While current stationary FC measurements lack sensitivity at the single-subject level, our investigation focuses on dynamic FC using resting-state functional MRI (rs-fMRI) and introduces the Generalized Auto-Regressive Conditional Heteroscedastic Dynamic Conditional Correlation (DCC-GARCH) model. Our findings demonstrate the superior sensitivity of DCC-GARCH to CSF *Aβ* status, and offer key insights into dynamic functional connectivity analysis in AD.

## 1 Introduction

Alzheimer’s disease (AD) is characterized by a prolonged latent phase, i.e. before noticeable symptoms manifest. This emphasizes the importance of developing sensitive biomarkers capable of identifying the preclinical stages of the disease. An accurate biomarker would facilitate the detection of AD in its early phases, enabling the implementation of disease-modifying therapies that might prevent cognitive impairment. [1].

In 2018, the National Institute on Aging and the Alzheimer’s Association (NIA-AA) introduced a ground-breaking research framework for AD, which fundamentally redefines the disease based on its underlying pathological processes, observable through biomarkers [2]. Unlike traditional diagnostic methods relying on clinical consequences or symptoms, this novel framework transforms the definition of AD in living individuals from a syndromal concept to a biological construct. According to this paradigm, the initiation of AD pathological changes in the brain begins with the formation of amyloid beta (*Aβ*) plaques, and subsequent presence of neurofibrillary tangles confirms the transition to AD. This biomarker-centered definition remains independent of clinical symptoms, indicating that biomarkers are poised to hold a central role in future clinical and research settings for AD.

Biomarkers capable of detecting the earliest pathological changes in the brain (i.e. pathological accumulation of *Aβ*) during the latent preclinical stage of AD offer a critical “window of opportunity.” During this stage, before irreversible damage occurs, therapeutic interventions may have their greatest impact. Currently, widely accepted biomarkers of *Aβ* deposition include cortical amyloid PET ligand binding or low levels of Cerebrospinal fluid (CSF), *Aβ*42. Moreover, studies have demonstrated that normalizing CSF *Aβ*42 concentration to the level of total *Aβ* using the *Aβ*42*/*40 ratio can enhance the differentiation between AD and controls [3, 4, 5, 6, 7]. Despite their significant success and invaluable contributions to research, these biomarkers may face challenges in population-based and clinical settings due to limited availability, cost, associated radiation exposure, and invasiveness.

More recent work indicates that soluble tau species (181-p-tau and 217-p-tau) also correlate well with significant *Aβ* deposition [8, 9]. However, despite the high cost, fluid markers of Alzheimer’s pathophysiology do not give a sense of the neurophysiological impact of the disease. Consequently, there is an urgent need for widely accessible, non-invasive AD biomarkers that can accurately detect the early stages of the physiologic disruptions of AD. Despite substantial efforts in this field, the development of such biomarkers remains a formidable challenge.

The identification of functional connectivity (FC) changes that correlate with AD pathology has suggested their potential use as non-invasive biomarkers for *Aβ* deposition [10, 11, 12, 13]. Reduced FC within the default mode network (DMN) has been observed in the early stages of AD [14, 15, 16, 17]. This reduction is associated with *Aβ* pathology [18, 19, 20, 21, 22, 23]. Resting-state functional MRI (rs-fMRI) can measure these changes non-invasively. However, despite encouraging results at the group level, current FC measurements do not yet have the sensitivity to act as standalone biomarkers [24].

The limited sensitivity of FC measurements may be partly attributed to the prevailing assumption of stationarity of FC in many studies. This assumption implies constant FC between brain regions throughout the entire scan duration (typically 5 to 10 minutes), represented by a single parameter derived from several minutes of data [25, 26]. However, evidence from electrophysiology suggests that FC undergoes dynamic changes within seconds to minutes [27, 28, 29, 30], exhibiting variations in both strength and direction [31, 32, 33, 34]. In the resting state, where mental activity is unrestricted, these dynamics may be even more pronounced.

The sliding-window approach has been a common approach for studying dynamic temporally-varying brain connectivities [34, 28, 31]. In this approach, a fixed-length window is employed to compute the correlation coefficient using data within the window. By shifting the window incrementally across time, a time-varying connectivity measurement between brain regions can be obtained. Although simple to implement, this approach has several important limitations. These include the utilization of arbitrary window lengths, the inability to handle abrupt changes, and the uniform weighting assigned to all observations within the selected window while disregarding older observations [35]. In particular, the choice of window length significantly influences the analysis results. Overly long window lengths obscure the true temporal dynamics in FC, diminishing the ability to capture essential information. Excessively short window lengths introduce inflated variance and may confound noise with genuine signals, potentially leading to misleading interpretations [36]. To address these limitations, we propose a novel method based on the Generalized Autoregressive Conditional Heteroscedastic Dynamic Conditional Correlation (DCC-GARCH) model [37, 38]. This model belongs to the volatility time series family, originally designed for financial applications but recently introduced to model dynamic brain connectivity networks in neuroscience studies [35, 39, 40]. The DCC-GARCH model provides a plausible mechanism to capture the temporal variation in brain networks, allowing easy interpretation of the model parameters. Specifically, the two model parameters *α* and *β* (to be explained in Section 2.2.2) offer valuable insights into the dynamic behavior of the FC system over time, reflecting its responsiveness to new stimuli and dependency on past brain states. We demonstrate that our method provides superior sensitivity compared to stationary FC measurement methods, enabling the detection of subtle pathological changes in AD and facilitating the development of non-invasive, MR-based AD biomarkers.

To meet the DCC-GARCH model’s assumptions, the time series observations for each node should exhibit no serial correlation after removing the mean signal (see Section 2.2.2). To achieve this, we apply a pre-whitening step as a common preprocessing approach to mitigate or eliminate serial correlations in time series observations. This procedure involves estimating the serial correlations from the observed data and utilizing the inverse correlation matrix for de-correlating the observations. While the literature on applying DCC-GARCH models to dynamic brain networks lacks relevant discussions on pre-whitening, the broader fMRI analysis literature extensively discusses appropriate approaches for pre-whitening [41, 42, 43, 44, 45, 46]. Traditional approaches, such as the autoregressive moving average (ARMA) model with fixed order in AFNI [47] or the global AR(1) model in SPM [48], often leave significant residual serial correlations in the processed data [49]. Recent studies propose using higher-order ARMA models with spatially-varying coefficients [45, 50] that demonstrate effective pre-whitening. For our study, we adopt the IDAR approach, which is an iterative, data-driven pre-whitening procedure based on AR models [49], ensuring improved performance in handling serial correlations and providing more reliable estimates for subsequent DCC-GARCH modeling.

## 2 Modeling Dynamic Functional Connectivity

Figure 1 illustrates the details of the analysis procedures. The collected rs-fMRI data undergoes standard preprocessing. Next, we compute time series observations for each node in the default mode network (DMN). These observations are obtained using a brain mask created through group independent component analysis (Group ICA) (FSL MELODIC, [51]). Then, the DCC-GARCH model quantifies the dynamic functional connectivity for each subject. Finally, we conduct a group comparison analysis to investigate the differences in dynamic connectivity patterns between AD patients and control subjects.

**Figure 1:**
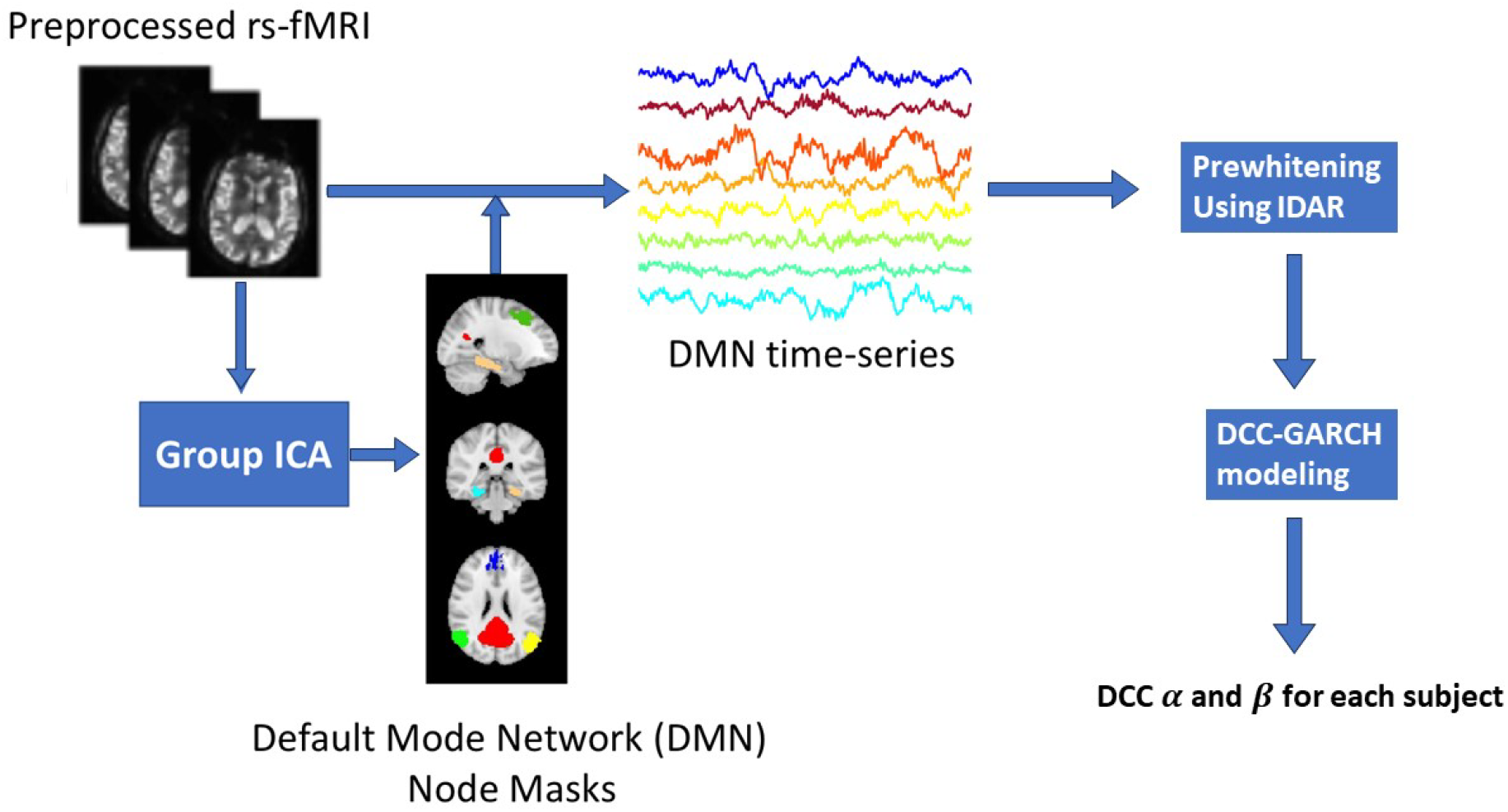
Flow chart of the analysis procedure.

### 2.1 Data

In this study, we included 87 subjects (mean age = 74 yr, range = 54 to 91 yr). All participants enrolled in UW ADRC (NIH P50 AG005136) Clinical Core, and were recruited with either normal cognition or mild cognitive impairment. All MRI data were acquired using a research-dedicated 3T Philips Achieva scanner equipped with a 32-channel receiver coil. During the scanning session, subjects were instructed to lie in the scanner with their eyes open while wearing a Pearl-Tec^®^ Crania to minimize head motion.

For registration purposes and volumetric measurements, we acquired 3D T1 data from all participants. The 3D structural images were obtained using a conventional MPRAGE sequence with the following parameters: spatial resolution = 1 *×* 1 *×* 1 mm^3^, 150 slices, flip angle = 8°, TR/TE = 8.8/4.6 ms, SENSE acceleration factor = 2, and matrix size = 256 *×* 256 *×* 176.

Rs-fMRI data was collected using axial whole-brain multi-echo T2-weighted sequence over a 10-minute scan period. These images used 3.5 mm isotropic voxels and had TR/TEs of 2,500/9.5, 27.5, and 45.5 ms. Following this, we carried out motion correction using FSL’s MCFLIRT software [51]. Non-brain matter was then removed using the FSL Brain Extraction Tool [51]. The multi-echo BOLD data were then processed using AFNI’s specialized module TEDANA, for multi-echo planar imaging and analysis with independent component analysis (ME-ICA). TEDANA allows for the differentiation between BOLD (neuronal) and non-BOLD (artifact) components, leveraging the characteristic linear echo-time dependence of BOLD T2 signals [52]. This approach produced recombined images optimally weighted across the three echo times, along with ME-ICA-denoised time series and spatial component maps. Subsequently, the data were co-registered to T1 images using FSL. After that, high-pass temporal filtering was applied, and the global signal and motion parameters were regressed from the data.

CSF samples were collected via lumbar puncture from all subjects. Levels of *Aβ*42 and *Aβ*40 in CSF samples were measured using multiplex bead assays (Luminex xMAP) run on the Luminex 200 instrument according to the manufacturer’s instructions [53]. Based on their CSF *Aβ*42*/*40 ratio, we divided the subjects into two groups: *Aβ*+ and *Aβ*−, using the internally determined cut-off threshold of 0.11 [54]. 25 subjects with CSF *Aβ*42*/*40 *<* 0.11 are classified as *Aβ*+ AD patients, and the rest 62 subjects are *Aβ*− control subjects.

**Table 1:** Demographics of the subjects.

|  | AD (N=25) | Control (N=62) | Overall (N=87) |
| --- | --- | --- | --- |
| Age |  |  |  |
| Mean (SD) | 74.0 (8.00) | 73.5 (8.86) | 73.6 (8.58) |
| Median [Min, Max] | 74.0 [60.0, 87.0] | 74.5 [54.0, 91.0] | 74.0 [54.0, 91.0] |
| Gender |  |  |  |
| F | 14 (56.0%) | 36 (58.1%) | 50 (57.5%) |
| M | 11 (44.0%) | 26 (41.9%) | 37 (42.5%) |
| Ethnicity |  |  |  |
| NA | 0 (0%) | 1 (1.6%) | 1 (1.1%) |
| Hispanic | 0 (0%) | 1 (1.6%) | 1 (1.1%) |
| Non-Hispanic | 25 (100%) | 60 (96.8%) | 85 (97.7%) |

### 2.2 Group-level Analysis

For each FC network, we calculated two sets of FC measures from the rs-fMRI dataset: static network modeling approach using dual regression, and dynamic network modeling approach using DCC-GARCH. Next, we explain the two modeling approaches.

#### 2.2.1 Static Network: Dual Regression

We measured the static strength of the DMN via the “dual regression” approach [55]. We first identified the standard space maps of the group average DMN by independent component analysis (FSL MELODIC) of the group-concatenated time series data. For each subject, the group-average DMN map was then regressed into that subject’s 4D space-time dataset using spatial predictors in a multiple regression, yielding a set of subject-specific time series reflecting DMN activity levels. These time series were then regressed back into the same 4D dataset using temporal predictors in another multiple regression, leading to subject-specific DMN maps. For each subject, we computed the average dual regression coefficient of all voxels within the DMN mask (mask obtained from group-average DMN map, thresholded at p *<* 0.001). This average served as the stationary FC measure of the DMN. Higher stationary FC measures imply a stronger average FC throughout the scan duration.

#### 2.2.2 Dynamic network: DCC-GARCH

We represented the FC of DMN via the conditional correlations among DMN nodes, and applied DCC-GARCH model to describe its temporal variation. The following 8 DMN nodes were considered: left and right superior frontal gyrus (SFG), medial prefrontal cortex (MPFC), posterior cingulate cortex (PCC), left and right parahippocampal cortex (PHC), left and right posterior inferior parietal lobule (pIPL). We extracted the corresponding time-series for each of the 8 nodes by averaging the measured signals over the mask voxels. Before performing the dynamic network analysis, we applied IDAR-based prewhitening procedure to each time series to remove the serial correlations [49].

We first explain the formulation of the DCC-GARCH model. Suppose we have a multivariate time series 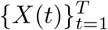 with *X*(*t*) ∈ μ^*m*^. The time series can be decomposed into a mean component *μ*(*t*) and a residual component *ϵ*(*t*):

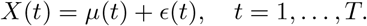

The mean process *μ*(*t*) can be modeled as a constant or with a time series model. Here we use a constant mean component. The residual component *ϵ*(*t*) is modeled by 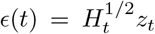, where *z*_*t*_ is a white noise process with variance *I*_*m*_. The conditional covariance matrix *H*_*t*_ conditions on the past information up to time *t*− 1. The residual component model indicates *ϵ*(*t*) is serially uncorrelated, even though it is still serially dependent.

The DCC-GARCH model assumes

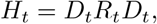

where the dynamic conditional covariance matrix 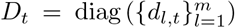 is a diagonal matrix with time-varying standard deviations *d*_*l,t*_ along its diagonal, and *R*_*t*_ is the dynamic conditional correlation matrix that represents the dynamic FC among brain nodes. The dynamic standard deviations *d*_*l,t*_ are usually modeled by a GARCH(1,1) process:

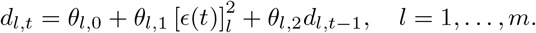

Higher order GARCH models are also applicable for the dynamic standard deviations, but the GARCH(1,1) model is usually adequate [56, 39].

The dynamic correlation matrix is modeled by

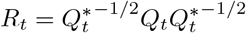

where 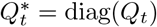 is the diagonal matrix with the diagonal elements of *Q*_*t*_, and

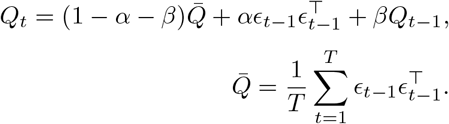

This formulation ensures a proper correlation matrix *R*_*t*_ that is positive-definite with diagonal entries equal to 1. The two parameters, *α* and *β*, characterize the temporal dynamics in the correlation network: *α* represents how the correlation matrix dynamically varies as a response to new stimulates, and *β* quantifies the degree of dependence of the correlations on their past values. In Figure 2, we provide a visualization illustrating the dynamic profile of correlations in a simulated 3-node network, highlighting the impact of *α* and *β* values. To generate the data from a DCC-GARCH model, we employed the R function simulateDCC from the ccgarch2 package (V0.0.0-42, [57]). For the simulations shown in Figure 2, we set the residual covariance matrix as

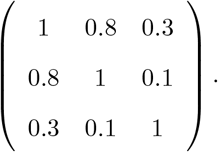

**Figure 2:**
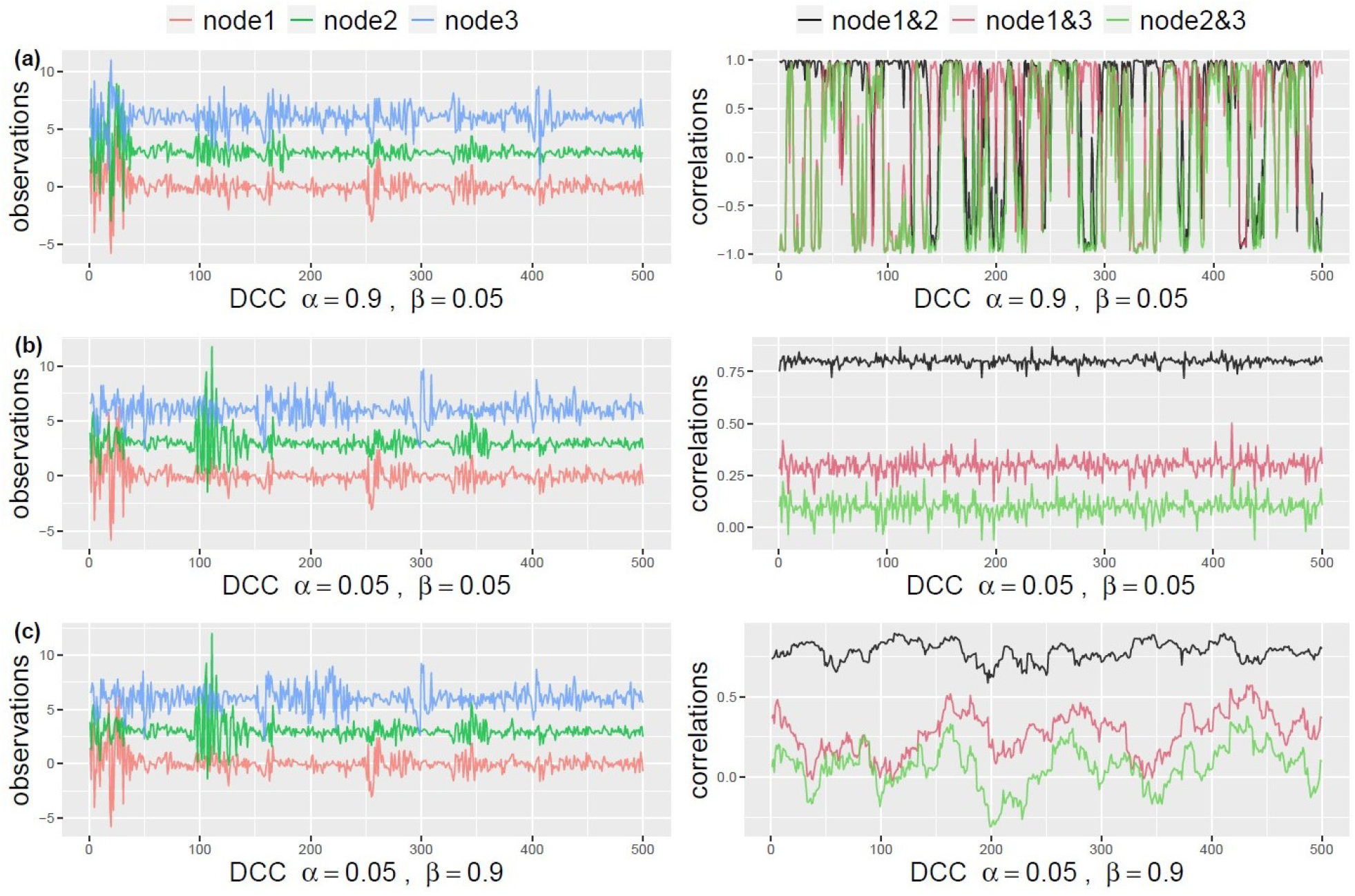
Illustration of the simulated dynamic profile of correlations with respect to different *α* and *β* values. We simulate three sets of time series observations for a 3-node network based on DCC-GARCH model. Each set has different parameter values for (*α, β*). The left side plots show the simulated signals from three nodes, and the right side plots show the dynamic profile of the corresponding pairwise correlations.

Additionally, for the GARCH(1,1) model of the dynamic standard deviations, we set all the coefficient parameters to 0.5. As shown in the plots, both *α* and *β* being small leads to stable correlations that resemble white noise (Figure 2b). Conversely, when *α* is small and *β* is large, the correlation pattern becomes “sticky”, changing slowly over time (Figure 2c). On the other hand, large values of *α* and small values of *β* generate rapid changes in correlations over time (Figure 2a). It is crucial to highlight that the dynamic profile of the correlations cannot be identified through simple visual examination of the observed signals. This underscores the necessity of employing appropriate models to accurately capture the temporal dynamics.

To analyze the dynamic correlation profiles from the fMRI data, the DCC-GARCH model was estimated via maximizing the quasi-likelihood, and implemented using the R package rmgarch [58]. After obtaining the subject-level estimates for the dynamic characterization parameters *α* and *β*, we collected the estimates and conduct a two-sample t-test for a group-level comparison between *Aβ*+ AD patients and *Aβ*− control subjects. We tested for group differences at 0.05 significance level.

## 3 Results

We begin by visualizing the dynamic correlation profile of an example subject based on the raw and whitened fMRI data using the sliding window approach. In this approach, we employed a simple rectangular window function. For the purpose of exposition, we focused on visualizing the correlations between specific pairs of brain nodes, namely PCC and SFG left, PCC and MPFC, PCC and PHC left, and PCC and pIPL left. As suggested in [59], the choice of a suitable minimum window length is influenced by the minimum frequency of the fMRI measurement. In our dataset, a window length of 100 seconds (equivalent to 50 data points) seemed appropriate. For comparison, we also included window lengths of 50s, 200s, and 400s. As depicted in Figure 3, the observed correlations exhibited temporal variability. The variability could get substantial, resulting in correlations that not only change sign, but also exhibited a large amplitude that can be more than twice the magnitude of the static correlation measurement. However, the magnitude of this variation was highly dependent on the window length, making it challenging to distinguish between genuine underlying dynamics and spurious fluctuations. Comparing the correlation profiles estimated from raw and whitened data, we observed that the dynamic pattern was not significantly altered by the whitening procedure. However, there was a notable decrease in the correlation between PCC and MPFC for this subject, with the static correlation measurement decreasing from 0.51 to 0.33.

**Figure 3:**
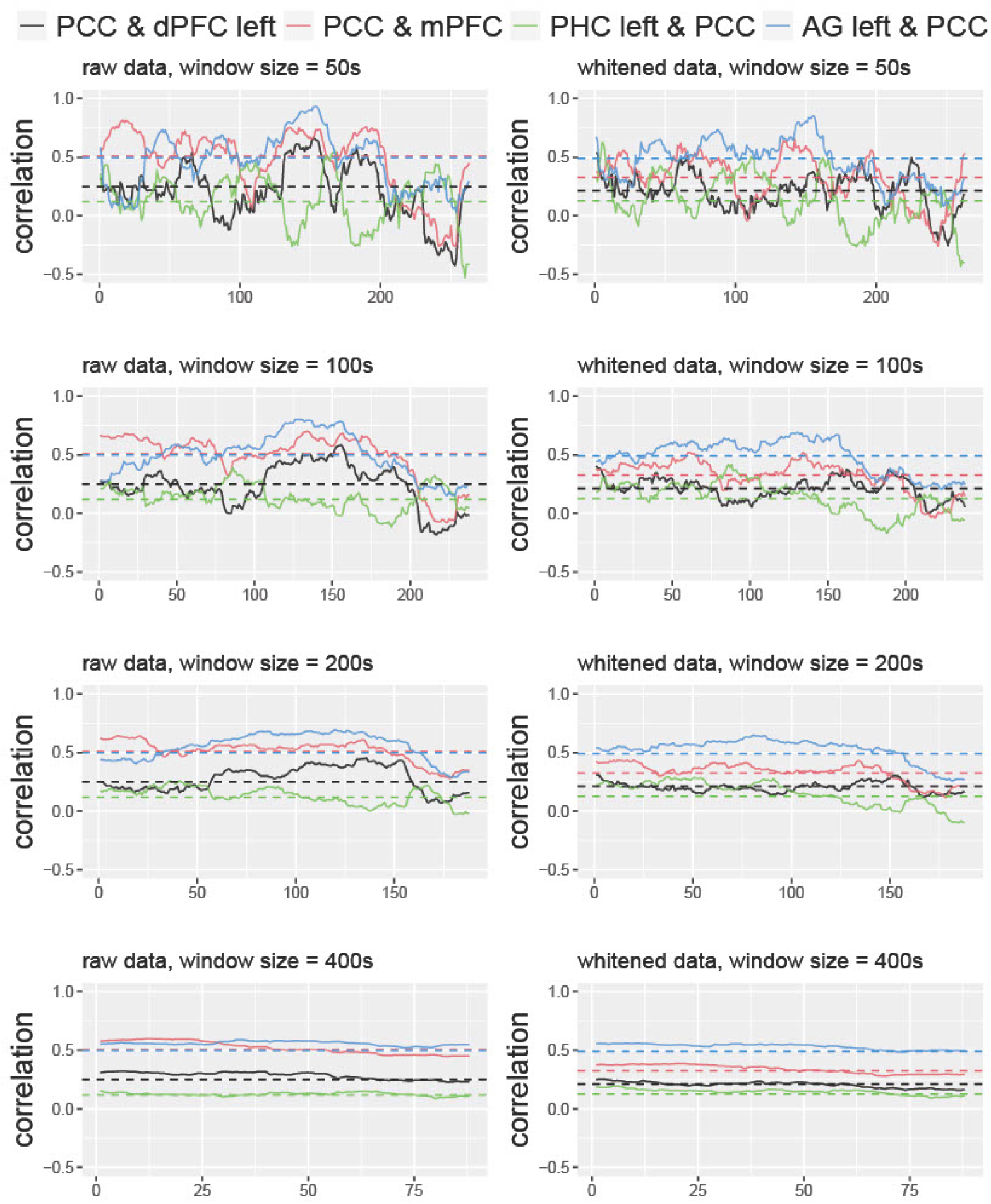
The dynamic correlation profile obtained via the sliding window approach, using the raw data and the whitened data, for an example subject. We visualize the correlation profile of four chosen brain node pairs. Correlations are computed using the rectangular window function at window length = 50s, 100s, 200s, 400s. The dashed lines represent the standard static correlation coefficient for each node pair, computed using data from the full time course.

Figure 4 compares the sliding-window-based correlations with the correlation profiles based on the estimated DCC-GARCH model. Three example subjects whose correlation profiles vary are shown. The correlations calculated with the DCC-GARCH model showe more noticeable differences among the subjects. Notably, the estimated correlation trajectories capture certain variation patterns identified by the sliding window approach. For instance, in subject 1, large drops in the correlations between PCC and SFG left were observed towards the end of the observation period.

**Figure 4:**
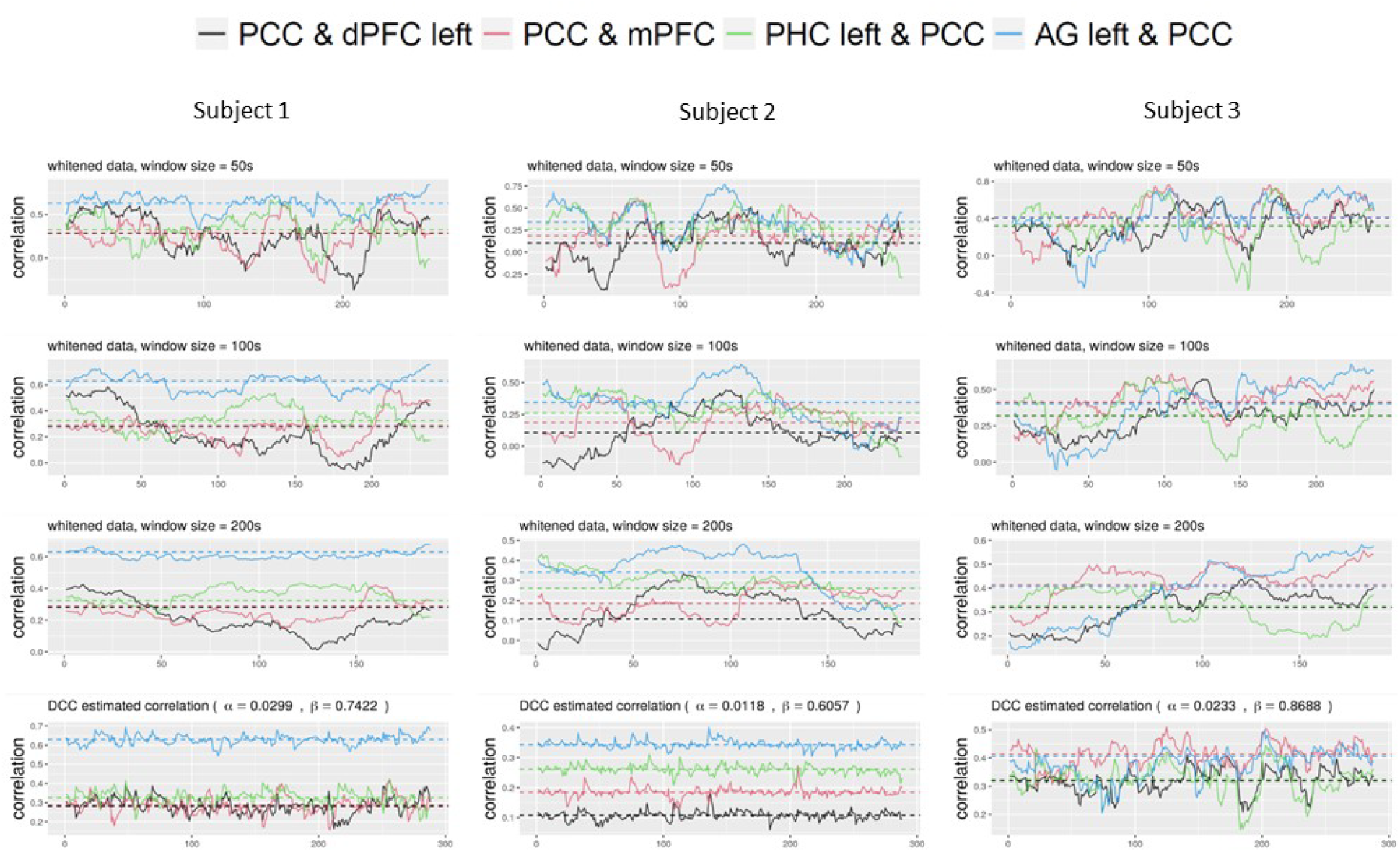
The dynamic correlation profile of three example subjects, using the whitened data, based on the sliding window approaches (top three rows) and the DCC-GARCH approach (last row). The sliding window approach applies the rectangular window function at window length = 50s, 100s, 200s. We visualize the correlation profile of four chosen brain node pairs. The dashed lines represent the standard static correlation coefficient for each node pair, computed using data from the full time course.

By fitting separate DCC-GARCH models to each subject in the dataset, we observed significant group differences at 0.05 level in terms of the subject-level dynamic characterization parameter *α* between the healthy controls and the AD subjects (Figure 5). The group difference in dynamic characterization parameter *β* was marginally significant at 0.1 level. The AD subjects in general had lower *α* values and higher *β* values, indicating less responsiveness to outside stimuli and more dependence on the previous brain’s connectivity status in the DMN compared to the healthy subjects. In contrast, the stationary measurement based on standard dual-regression ICA analysis (Figure 5) did not show significant group difference.

**Figure 5:**
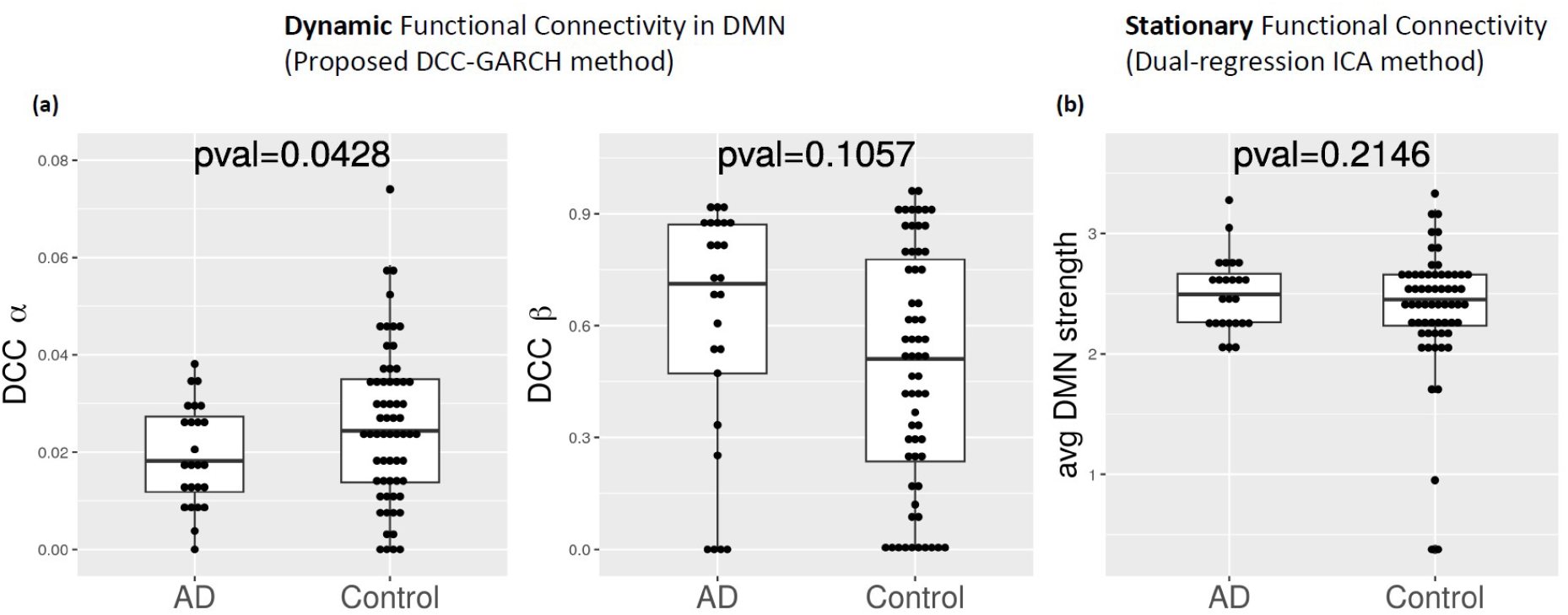
(a) Boxplots of estimated DCC-GARCH *α* and *β* parameters calculated using the proposed method for all subjects. Each dot represents the parameter estimate for a single subject. Subjects were divided into two groups based on their CSF *Aβ* profile: 1) AD patients (CSF *Aβ*42/ *Aβ*40 *<* 0.11), and 2) control (CSF *Aβ*42/ *Aβ*40 ≥ 0.11). p-values of two-sample t-test for group differences in parameter estimates are reported. (b) Boxplots of average stationary functional connectivity in DMN, calculated using dual-regression ICA analysis.

We then assessed the sensitivity and specificity of the estimated DCC-GARCH parameters in identifying AD patients (Figure 6). To classify a subject as AD, we employed varying threshold values for the estimated DCC-GARCH parameters *α* and/or *β*. With the thresholds ranging from 0 to 1, we generated ROC curves and calculated their corresponding AUC. In the “combined” version where both *α* and *β* values were used for classification, we employed the Support Vector Machine (SVM) approach with the radial kernel, where the tuning parameters (*γ* and regularization constant *C*) were selected to maximize AUC based on 10-fold cross-validation. When used separately for classification, the *α* and *β* parameters had moderate power to identify AD patients, each with an AUC of 0.6. We observed higher discriminative power for patient diagnosis when using both *α* and *β* values for classification, yielding an AUC of 0.7. On the contrary, the stationary functional connectivity measurement of average DMN strength computed from dual-regression ICA analysis shows no classification power for patient diagnosis.

**Figure 6:**
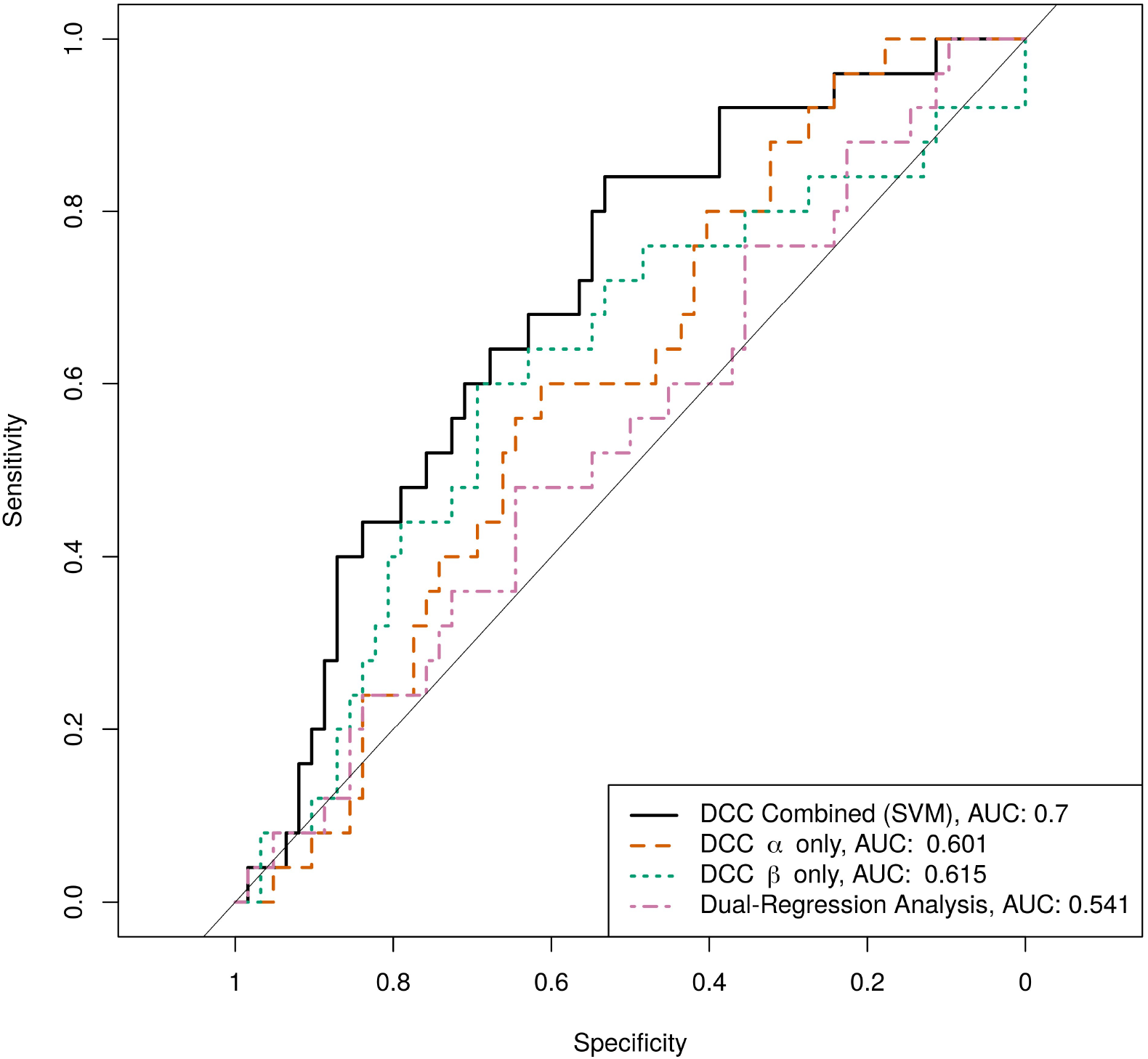
ROC curves and AUC for identifying AD patients based on estimated DCC-GARCH parameters. The ‘DCC *α* only’ and ‘DCC *β* only’ curves are based on thresholding the estimated *α* or *β* values separately, the ‘DCC Combined’ ROC curve is yielded from using *α* and *β* values via the SVM approach. The ‘Dual-Regression Analysis’ ROC curve is based on average DMN strength computed from dual-regression ICA analysis.

## 4 Discussion

Our study sheds light on the importance of considering dynamic network configurations and the limitations of stationary measures of functional connectivity when searching for biomarkers for Alzheimer’s Disease. To capture the temporal dynamics of brain network connectivity, we proposed using a DCC-GARCH model. Proper pre-processing steps, especially an effective pre-whitening procedure to eliminate serial correlations, are necessary for the validity of applying the DCC-GARCH model, which ensures the model assumptions are met. By addressing these key issues, our findings contribute to the understanding of functional connectivity analysis in AD research and offer insights into the development of non-invasive and widely available biomarkers for the early detection of the disease. The results of our study indicate that the proposed DCC-GARCH modeling approach for dynamic functional connectivity in the default mode network (DMN) is sensitive to the CSF *Aβ* status. Specifically, the FC measurements of AD patients seem to exhibit higher lagging dependency and reduced responsiveness to new information compared to healthy controls. The resulting DCC-GARCH model parameters show reasonable power in identifying AD patients in the population, indicating its potential as a sensitive biomarker for early detection of AD based on *Aβ* deposition. We anticipate that with a larger number of subjects, the differences between the two groups would become more significant. Additionally, we expect that increased sample size would lead to better predictability of our model.

It is important to note that the DCC-GARCH model provides a plausible description of the temporal dynamics in the DMN, considering the data from the perspective of autoregressive dependency. While it may not represent the true or best model for describing the brain mechanism, it effectively captures important variation patterns identified by the sliding-window-based approach. The estimated dynamic functional correlation profiles using the DCC-GARCH model tend to be more stable and more concentrated around the static correlation measurement compared to the sliding-window-based approach, reflected in smaller standard deviations over the time course. Further investigation should explore alternative perspectives, such as network topology or identifying change points in correlation profiles, to gain deeper insights into potential group differences. By examining multiple aspects, we can enhance our understanding of the dynamic functional connectivity alterations in AD and uncover more sensitive biomarkers for early detection.

In summary, our study provides valuable insights into the analysis of dynamic functional connectivity in AD and the potential for developing biomarkers for the disease. The utilization of the DCC-GARCH model offers a promising approach to capturing the temporal dynamics of brain network connectivity. Future research should focus on validating our findings in larger cohorts of independent subjects and exploring additional factors that may differentiate the dynamic correlation profiles observed between AD patients and healthy individuals. By continuing to advance our understanding of functional connectivity dynamics, we can pave the way for improved diagnosis and monitoring of AD, ultimately leading to more effective interventions and treatments.

## 5 Data Availability

Researchers can obtain de-identified raw imaging data by accessing the National Alzheimer’s Coordinating Center (NACC) website, which includes DICOM format raw imaging files and variables from the NACC Uniform Dataset Version 3.0. Additionally, investigators with an IRB-approved study and a designated UW ADRC collaborator may be granted access to the UW Alzheimer’s Disease Research Center (ADRC) Imaging and Biomarker Core database. The imaging data utilized in this study is available in BIDS format and is stored on an XNAT platform in the UW Integrated Brain Imaging Center. For inquiries, please reach out to Dr. Thomas Grabowski MD, Lead of the Imaging and Biomarker Core at UW ADRC.

## Notes

### Competing Interest Statement

The authors have declared no competing interest.

## References

[1] Dennis J Selkoe. Presenilin, notch, and the genesis and treatment of alzheimer’s disease. Proceedings of the National Academy of Sciences, 98(20):11039–11041, 2001.

[2] Clifford R Jack Jr, David A Bennett, Kaj Blennow, Maria C Carrillo, Billy Dunn, Samantha Budd Haeberlein, David M Holtzman, William Jagust, Frank Jessen, Jason Karlawish, et al. Nia-aa research framework: toward a biological definition of alzheimer’s disease. Alzheimer’s & Dementia, 14(4):535–562, 2018.

[3] Inês Baldeiras, Isabel Santana, Maria João Leitão, Maria Helena Ribeiro, Rui Pascoal, Diana Duro, Raquel Lemos, Beatriz Santiago, Maria Rosário Almeida, and Catarina Resende Oliveira. Cerebrospinal fluid aβ40 is similarly reduced in patients with frontotemporal lobar degeneration and alzheimer’s disease. Journal of the neurological sciences, 358(1-2):308–316, 2015.

[4] Piotr Lewczuk, Natalia Lelental, Philipp Spitzer, Juan Manuel Maler, and Johannes Kornhuber. Amyloid-β 42/40 cerebrospinal fluid concentration ratio in the diagnostics of alzheimer’s disease: validation of two novel assays. Journal of Alzheimer’s disease, 43(1):183–191, 2015.

[5] Piotr Lewczuk, Hermann Esselmann, Markus Otto, Juan Manuel Maler, Andreas Wolfram Henkel, Maria Kerstin Henkel, Oliver Eikenberg, Christof Antz, Wolf-Rainer Krause, Udo Reulbach, et al. Neurochemical diagnosis of alzheimer’s dementia by csf aβ42, aβ42/aβ40 ratio and total tau. Neurobiology of aging, 25(3):273–281, 2004.

[6] Shorena Janelidze, Henrik Zetterberg, Niklas Mattsson, Sebastian Palmqvist, Hugo Vanderstichele, Olof Lindberg, Danielle van Westen, Erik Stomrud, Lennart Minthon, Kaj Blennow, et al. Csf aβ42/aβ40 and aβ42/aβ38 ratios: better diagnostic markers of alzheimer disease. Annals of clinical and translational neurology, 3(3):154–165, 2016.

[7] Inês Baldeiras, Isabel Santana, Maria João Leitão, Helena Gens, Rui Pascoal, Miguel Tábuas-Pereira, José Beato-Coelho, Diana Duro, Maria Rosário Almeida, and Catarina Resende Oliveira. Addition of the aβ42/40 ratio to the cerebrospinal fluid biomarker profile increases the predictive value for underlying alzheimer’s disease dementia in mild cognitive impairment. Alzheimer’s research & therapy, 10(1):1–15, 2018.

[8] Nicolas R Barthélemy, Kanta Horie, Chihiro Sato, and Randall J Bateman. Blood plasma phosphorylated-tau isoforms track cns change in alzheimer’s disease. Journal of Experimental Medicine, 217(11):e20200861, 2020.

[9] Michelle M Mielke, Clinton E Hagen, Jing Xu, Xiyun Chai, Prashanthi Vemuri, Val J Lowe, David C Airey, David S Knopman, Rosebud O Roberts, Mary M Machulda, et al. Plasma phospho-tau181 increases with alzheimer’s disease clinical severity and is associated with tau-and amyloid-positron emission tomography. Alzheimer’s & Dementia, 14(8):989–997, 2018.

[10] Eek-Sung Lee, Kwangsun Yoo, Young-Beom Lee, Jinyong Chung, Ji-Eun Lim, Bora Yoon, and Yong Jeong. Default mode network functional connectivity in early and late mild cognitive impairment. Alzheimer Disease & Associated Disorders, 30(4):289–296, 2016.

[11] Kamil A Grajski, Steven L Bressler, Alzheimer’s Disease Neuroimaging Initiative, et al. Differential medial temporal lobe and default-mode network functional connectivity and morphometric changes in alzheimer’s disease. NeuroImage: Clinical, 23:101860, 2019.

[12] Jessica S Damoiseaux, Katherine E Prater, Bruce L Miller, and Michael D Greicius. Functional connectivity tracks clinical deterioration in alzheimer’s disease. Neurobiology of aging, 33(4):828–e19, 2012.

[13] Marcio Luiz Figueredo Balthazar, Brunno Machado de Campos, Alexandre Rosa Franco, Benito Pereira Damasceno, and Fernando Cendes. Whole cortical and default mode network mean functional connectivity as potential biomarkers for mild alzheimer’s disease. Psychiatry Research: Neuroimaging, 221(1):37–42, 2014.

[14] Michael D Greicius, Gaurav Srivastava, Allan L Reiss, and Vinod Menon. Default-mode network activity distinguishes alzheimer’s disease from healthy aging: evidence from functional mri. Proceedings of the National Academy of Sciences, 101(13):4637–4642, 2004.

[15] Christian Sorg, Valentin Riedl, Mark Muhlau, Vince D Calhoun, Tom Eichele, Leonhard Läer, Alexander Drzezga, Hans Förstl, Alexander Kurz, Claus Zimmer, et al. Selective changes of resting-state networks in individuals at risk for alzheimer’s disease. Proceedings of the National Academy of Sciences, 104(47):18760–18765, 2007.

[16] Liang Wang, Yufeng Zang, Yong He, Meng Liang, Xinqing Zhang, Lixia Tian, Tao Wu, Tianzi Jiang, and Kuncheng Li. Changes in hippocampal connectivity in the early stages of alzheimer’s disease: evidence from resting state fmri. Neuroimage, 31(2):496–504, 2006.

[17] Hong-Ying Zhang, Shi-Jie Wang, Jiong Xing, Bin Liu, Zhan-Long Ma, Ming Yang, Zhi-Jun Zhang, and Gao-Jun Teng. Detection of pcc functional connectivity characteristics in resting-state fmri in mild alzheimer’s disease. Behavioural brain research, 197(1):103–108, 2009.

[18] Willem Huijbers, Elizabeth C Mormino, Sarah E Wigman, Andrew M Ward, Patrizia Vannini, Donald G McLaren, J Alex Becker, Aaron P Schultz, Trey Hedden, Keith A Johnson, et al. Amyloid deposition is linked to aberrant entorhinal activity among cognitively normal older adults. Journal of Neuroscience, 34(15):5200–5210, 2014.

[19] Cindy Lustig, Abraham Z Snyder, Mehul Bhakta, Katherine C O’Brien, Mark McAvoy, Marcus E Raichle, John C Morris, and Randy L Buckner. Functional deactivations: change with age and dementia of the alzheimer type. Proceedings of the National Academy of Sciences, 100(24):14504–14509, 2003.

[20] Randy L Buckner, Abraham Z Snyder, Benjamin J Shannon, Gina LaRossa, Rimmon Sachs, Anthony F Fotenos, Yvette I Sheline, William E Klunk, Chester A Mathis, John C Morris, et al. Molecular, structural, and functional characterization of alzheimer’s disease: evidence for a relationship between default activity, amyloid, and memory. Journal of neuroscience, 25(34):7709–7717, 2005.

[21] Trey Hedden, Koene RA Van Dijk, J Alex Becker, Angel Mehta, Reisa A Sperling, Keith A Johnson, and Randy L Buckner. Disruption of functional connectivity in clinically normal older adults harboring amyloid burden. Journal of Neuroscience, 29(40):12686–12694, 2009.

[22] Yvette I Sheline, Marcus E Raichle, Abraham Z Snyder, John C Morris, Denise Head, Suzhi Wang, and Mark A Mintun. Amyloid plaques disrupt resting state default mode network connectivity in cognitively normal elderly. Biological psychiatry, 67(6):584–587, 2010.

[23] Elizabeth C Mormino, Andre Smiljic, Amynta O Hayenga, Susan H. Onami, Michael D Greicius, Gil D Rabinovici, Mustafa Janabi, Suzanne L Baker, Irene V. Yen, Cindee M Madison, et al. Relationships between beta-amyloid and functional connectivity in different components of the default mode network in aging. Cerebral cortex, 21(10):2399–2407, 2011.

[24] Stefan J Teipel, Coraline D Metzger, Frederic Brosseron, Katharina Buerger, Katharina Brueggen, Cihan Catak, Dominik Diesing, Laura Dobisch, Klaus Fliebach, Christiana Franke, et al. Multicenter resting state functional connectivity in prodromal and dementia stages of alzheimer’s disease. Journal of Alzheimer’s Disease, 64(3):801–813, 2018.

[25] Bharat Biswal, F Zerrin Yetkin, Victor M Haughton, and James S Hyde. Functional connectivity in the motor cortex of resting human brain using echo-planar mri. Magnetic resonance in medicine, 34(4):537–541, 1995.

[26] Zarrar Shehzad, AM Clare Kelly, Philip T Reiss, Dylan G Gee, Kristin Gotimer, Lucina Q Uddin, Sang Han Lee, Daniel S Margulies, Amy Krain Roy, Bharat B Biswal, et al. The resting brain: unconstrained yet reliable. Cerebral cortex, 19(10):2209–2229, 2009.

[27] R Matthew Hutchison, Thilo Womelsdorf, Elena A Allen, Peter A Bandettini, Vince D Calhoun, Maurizio Corbetta, Stefania Della Penna, Jeff H Duyn, Gary H Glover, Javier Gonzalez-Castillo, et al. Dynamic functional connectivity: promise, issues, and interpretations. Neuroimage, 80:360–378, 2013.

[28] Catie Chang and Gary H Glover. Time–frequency dynamics of resting-state brain connectivity measured with fmri. Neuroimage, 50(1):81–98, 2010.

[29] Enzo Tagliazucchi, Frederic Von Wegner, Astrid Morzelewski, Verena Brodbeck, and Helmut Laufs. Dynamic bold functional connectivity in humans and its electrophysiological correlates. Frontiers in human neuroscience, 6:339, 2012.

[30] George C O’Neill, Prejaas K Tewarie, Giles L Colclough, Lauren E Gascoyne, Benjamin AE Hunt, Peter G Morris, Mark W Woolrich, and Matthew J Brookes. Measurement of dynamic task related functional networks using meg. NeuroImage, 146:667–678, 2017.

[31] Daniel A Handwerker, Vinai Roopchansingh, Javier Gonzalez-Castillo, and Peter A Bandettini. Periodic changes in fmri connectivity. Neuroimage, 63(3):1712–1719, 2012.

[32] Ünal Sakoğlu, Godfrey D Pearlson, Kent A Kiehl, Y Michelle Wang, Andrew M Michael, and Vince D Calhoun. A method for evaluating dynamic functional network connectivity and task-modulation: application to schizophrenia. Magnetic Resonance Materials in Physics, Biology and Medicine, 23:351–366, 2010.

[33] David T Jones, Prashanthi Vemuri, Matthew C Murphy, Jeffrey L Gunter, Matthew L Senjem, Mary M Machulda, Scott A Przybelski, Brian E Gregg, Kejal Kantarci, David S Knopman, et al. Non-stationarity in the “resting brain’s” modular architecture. PloS one, 7(6):e39731, 2012.

[34] Elena A Allen, Eswar Damaraju, Sergey M Plis, Erik B Erhardt, Tom Eichele, and Vince D Calhoun. Tracking whole-brain connectivity dynamics in the resting state. Cerebral cortex, 24(3):663–676, 2014.

[35] Martin A Lindquist, Yuting Xu, Mary Beth Nebel, and Brain S Caffo. Evaluating dynamic bivariate correlations in resting-state fmri: a comparison study and a new approach. NeuroImage, 101:531–546, 2014.

[36] Hamed Honari, Ann S Choe, James J Pekar, and Martin A Lindquist. Investigating the impact of autocorrelation on time-varying connectivity. NeuroImage, 197:37–48, 2019.

[37] Robert F Engle III and Kevin Sheppard. Theoretical and empirical properties of dynamic conditional correlation multivariate garch, 2001.

[38] Robert Engle. Dynamic conditional correlation: A simple class of multivariate generalized autoregressive conditional heteroskedasticity models. Journal of Business & Economic Statistics, 20(3):339–350, 2002.

[39] R Riccelli, Luca Passamonti, Andrea Duggento, Maria Guerrisi, Iole Indovina, A Terracciano, and Nicola Toschi. Dynamical brain connectivity estimation using garch models: An application to personality neuroscience. In 2017 39th Annual International Conference of the IEEE Engineering in Medicine and Biology Society (EMBC), pages 3305–3308. IEEE, 2017.

[40] Namgil Lee and Jong-Min Kim. Dynamic functional connectivity analysis based on time-varying partial correlation with a copula-dcc-garch model. Neuroscience Research, 169:27–39, 2021.

[41] Wiktor Olszowy, John Aston, Catarina Rua, and Guy B Williams. Accurate autocorrelation modeling substantially improves fmri reliability. Nature communications, 10(1):1220, 2019.

[42] Brian Lenoski, Leslie C Baxter, Lina J Karam, José Maisog and Josef Debbins. On the performance of autocorrelation estimation algorithms for fmri analysis. IEEE Journal of Selected Topics in Signal Processing, 2(6):828–838, 2008.

[43] Molly G Bright, Christopher R Tench, and Kevin Murphy. Potential pitfalls when denoising resting state fmri data using nuisance regression. Neuroimage, 154:159–168, 2017.

[44] Mark W Woolrich, Brian D Ripley, Michael Brady, and Stephen M Smith. Temporal autocorrelation in univariate linear modeling of fmri data. Neuroimage, 14(6):1370–1386, 2001.

[45] Qingfei Luo, Masaya Misaki, Ben Mulyana, Chung-Ki Wong, and Jerzy Bodurka. Improved autoregressive model for correction of noise serial correlation in fast fmri. Magnetic resonance in medicine, 84(3):1293–1305, 2020.

[46] Nadège Corbin, Nick Todd, Karl J Friston, and Martina F Callaghan. Accurate modeling of temporal correlations in rapidly sampled fmri time series. Human brain mapping, 39(10):3884–3897, 2018.

[47] Robert W Cox. Afni: software for analysis and visualization of functional magnetic resonance neuroimages. Computers and Biomedical research, 29(3):162–173, 1996.

[48] William D Penny, Karl J Friston, John T Ashburner, Stefan J Kiebel, and Thomas E Nichols. Statistical parametric mapping: the analysis of functional brain images. Elsevier, 2011.

[49] Kun Yue, Jason M Webster, Thomas Grabowski, Ali Shojaei, and Hesamoddin Jahanian. Iterative data-adaptive autoregressive (idar) whitening procedure for long and short tr fmri. Manuscript submitted for publication., 2023.

[50] Ashish Kaul Sahib, Klaus Mathiak, Michael Erb, Adham Elshahabi, Silke Klamer, Klaus Scheffler, Niels K Focke, and Thomas Ethofer. Effect of temporal resolution and serial autocorrelations in event-related functional mri. Magnetic resonance in medicine, 76(6):1805–1813, 2016.

[51] Mark Jenkinson, Christian F Beckmann, Timothy EJ Behrens, Mark W Woolrich, and Stephen M Smith. Fsl. Neuroimage, 62(2):782–790, 2012.

[52] Prantik Kundu, Souheil J Inati, Jennifer W Evans, Wen-Ming Luh, and Peter A Bandettini. Differentiating bold and non-bold signals in fmri time series using multi-echo epi. Neuroimage, 60(3):1759–1770, 2012.

[53] Erika Oliveira Hansen, Natalia Silva Dias, Ivonne Carolina Bolanõs Burgos, Monica Vieira Costa, Andréa Teixeira Carvalho, Antonio Lucio Teixeira, Izabela Guimarães Barbosa, Lorena Aline Valu Santos, Daniela Valadão Freitas Rosa, Aloisio Joaquim Freitas Ribeiro, et al. Millipore xmap® luminex (hatmag-68k): An accurate and cost-effective method for evaluating alzheimer’s biomarkers in cerebrospinal fluid. Frontiers in Psychiatry, 12:716686, 2021.

[54] Orestes V Forlenza, Marcia Radanovic, Leda L Talib, Ivan Aprahamian, Breno S Diniz, Henrik Zetterberg, and Wagner F Gattaz. Cerebrospinal fluid biomarkers in alzheimer’s disease: diagnostic accuracy and prediction of dementia. Alzheimer’s & Dementia: Diagnosis, Assessment & Disease Monitoring, 1(4):455–463, 2015.

[55] Lisa D Nickerson, Stephen M Smith, Döst Öngür, and Christian F Beckmann. Using dual regression to investigate network shape and amplitude in functional connectivity analyses. Frontiers in neuroscience, 11:115, 2017.

[56] Elisabeth Orskaug. Multivariate dcc-garch model:-with various error distributions. Master’s thesis, Institutt for matematiske fag, 2009.

[57] Tomoaki Nakatani. ccgarch2: An R Package for Modelling Multivariate GARCH Models with Conditional Correlations, 2016. R package version 0.0.0-42, URL ¡http://CRAN.r-project/package=ccgarch¿.

[58] Alexios Galanos. rmgarch: Multivariate GARCH models., 2022. R package version 1. 3–9.

[59] Nora Leonardi and Dimitri Van De Ville. On spurious and real fluctuations of dynamic functional connectivity during rest. Neuroimage, 104:430–436, 2015.

